# Gaia: A Context-Aware Sequence Search and Discovery Tool for Microbial Proteins

**DOI:** 10.1101/2024.11.19.624387

**Authors:** Nishant Jha, Joshua Kravitz, Jacob West-Roberts, Antonio Camargo, Simon Roux, Andre Cornman, Yunha Hwang

## Abstract

Protein sequence similarity search is fundamental to genomics research, but current methods are typically not able to consider crucial genomic context information that can be indicative of protein function, especially in microbial systems. Here we present Gaia (Genomic AI Annotator), a sequence annotation platform that enables rapid, context-aware protein sequence search across genomic datasets. Gaia leverages gLM2, a mixed-modality genomic language model trained on both amino acid sequences and their genomic neighborhoods to generate embeddings that integrate sequence-structure-context information. This approach allows for the identification of functionally related genes that are found in conserved genomic contexts, which may be missed by traditional sequence- or structure-based search alone. Gaia enables real-time search of a curated database comprising over 85M protein clusters (defined at 90% sequence identity) from 131,744 microbial genomes. We compare the sequence, structure and context sensitivity of gLM2 embedding-based search against existing tools like MMseqs2 and Foldseek. We showcase Gaia-enabled discoveries of phage tail proteins and siderophore synthesis loci that were previously difficult to annotate with traditional tools. Gaia search is freely available at https://gaia.tatta.bio.

## 1 Introduction

Protein sequence similarity search is a foundational analytical technique in genomics research, critical for inferring protein function, predicting evolutionary relationships, and discovering novel genes. Tools like BLAST [10] and HMMER [14] have been widely used for these analyses, relying on sequence alignment methods to compare individual query sequences against databases of known proteins. While highly effective for detecting close homologs, these approaches can struggle to identify remote homologs that have diverged significantly.

Structural search methods are an effective alternative method to detect remote homology between proteins. Tools like Foldseek [51] perform structural similarity search on large databases of experimental and in silico predicted protein structures including the PDB [9] and AlphaFoldDB [52], and are capable of identifying highly similar structures with low sequence similarity to the provided input. However, relying on structural similarities results in limited insights for some intrinsically disordered proteins [3, 4], proteins with conformational changes [8], and proteins with low-confidence predicted structures [2]. Additionally, structural search requires pre-computed protein structures across large scale databases, incurring massive computational costs as protein databases grow to billion scale.

In recent years, the advent of biological language models has opened new avenues for sequence analysis. Models like ESM2 [27] and ESM3 [21] have shown promise in capturing complex patterns in protein sequences, enabling more sensitive remote homology detection. ESMFold, for example, provides accurate and scalable protein structure prediction, using only individual sequences as input. Tools like ProtTrek [47], pLM-BLAST [23], and PLMSearch [28] utilize protein language models for remote homology detection [16] by comparing against databases like SWISSProt [49] and ECOD [41]. Embedding-based search is highly efficient and scalable, showing similar or superior performance to alignment-based tools with increased sensitivity [28]. These approaches complement existing methods, offering alternative perspectives on protein relationships and functions.

However, a critical aspect often overlooked in current search methodologies is the genomic context of proteins, particularly informative of function in microbial genomics. Understanding the genomic neighborhood of a gene can provide crucial insights into its function and regulation. For example, the identification of CRISPR-associated genes relies on analyzing surrounding genomic regions [44]. Similarly, defense islands are identified by recognizing clusters of genes with shared annotations; ‘islands’ are defined as such by the co-location of these defense genes on the host genome [13, 29]. Traditionally, studies and analyses which utilize genome context are limited in throughput due to the slow, often manual, nature of the work required to inspect a full genomic locus and all the annotations within it.

## 2 Gaia

To address these challenges and bridge the gap between sequence, structure, and context-based analyses, we developed Gaia (Genomic AI Annotator). Gaia is an embedding-based search platform that enables fast and context-aware protein sequence similarity search over large genomic datasets.

Gaia enhances protein sequence analysis in genomic datasets by integrating contextual genomic information, a critical dimension often not considered in traditional search methods. The platform’s core is gLM2, a language model trained on the Open MetaGenome (OMG) dataset [12] (Fig. 1A). gLM2 is trained on both protein sequences and their surrounding genomic regions (including intergenic regions), generating features that capture complex relationships between genes and their genomic neighborhoods (up to 4096 tokens, corresponding to 9.7±3.3 genes). We fine-tune gLM2 (see Methods) yielding a structure-aligned and context-aware retrieval model (gLM2_embed) (Fig. 1B). gLM2’s context-awareness enables Gaia to identify functionally related proteins that may be missed by sequence-only or structure-only search, such as remote homologs in similar genomic contexts, or context-conserved multi-gene systems (i.e. defense islands and biosynthetic gene clusters).

**Figure 1:**
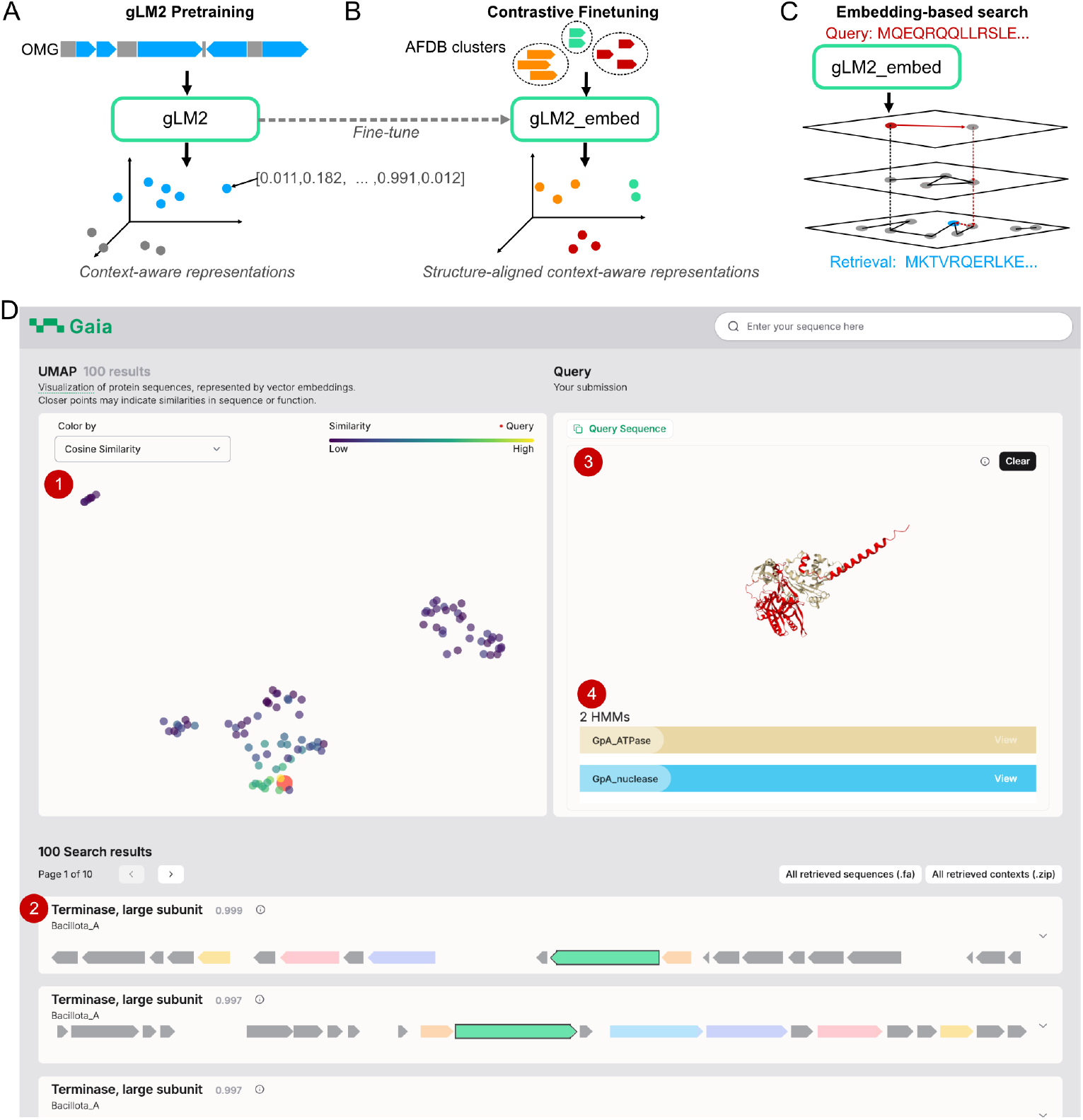
Gaia uses gLM2 representations to perform context-aware search. **(A)** gLM2 is a genomic language model trained on multi-gene metagenomic contigs from the Open MetaGenome (OMG) dataset, with a masked language modeling objective. gLM2 learns context-aware representations of proteins. **(B)** To improve retrieval of remote homologs, we fine-tune gLM2 representations to align them with structural clusters in AlphaFold database (AFDB) clusters, resulting in gLM2_embed. **(C)** Gaia’s embedding-based search uses the hierarchical navigable small world (HNSW) algorithm to rapidly find nearest neighbors in a protein vector database embedded by the gLM2_embed model. **(D)** Gaia search web server integrates (1) UMAP visualization of retrieved embeddings, (2) visualization of retrieved genomic contexts with top five most frequently co-occurring genes across the retrieved contigs, (3) ESMFold structure predictions of the query protein as well as all retrieved proteins and (4) detected Pfam HMMs. Additional information calculated and visualized by Gaia include local and global alignments, and sample metadata (Appendix A).

Gaia search takes in a single protein as a query, which is subsequently embedded into a gLM2_embed vector. We use the HNSW algorithm to conduct scalable nearest neighbor search using the cosine similarity metric (Fig. 1C) against a pre-computed vector database of 85M protein clusters (OG_prot_90, proteins from OpenGenome dataset clustered at 90% sequence identity). By default, Gaia search returns 100 nearest neighbors from OG_prot_90.

Gaia shifts the paradigm of sequence search from protein retrieval to genomic context retrieval. To facilitate validation and provide comprehensive insights, Gaia provides a clear visual representation of the genomic architectures of matched sequences with highlighted co-occurring genes, and incorporates results from additional tools such as BLASTp identity, PFAM HMM annotations [33], and ESMFold for protein structure prediction (Fig. 1D and Appendix A). This integrated approach allows users to efficiently retrieve and analyze relevant sequences, complete with functional annotations, structural predictions, and genomic context visualizations.

## 3 Results

We benchmark Gaia search against ESM2 embedding-based search, as well as other commonly used sequence and structural search tools (e.g. BLASTp, MMseqs2, Foldseek). We compare Gaia’s search sensitivities across three major axes of information pertaining to protein function: sequence (Fig. 2A), context (Fig. 2B) and structure (Fig. 2C). For sequence search sensitivity, we compare embedding-based search methods in their ability to retrieve the best non-self BLASTp matches of queries within top K nearest neighbors in the OG_prot_90 database. We find that Gaia search is significantly more sensitive when retrieving highly similar proteins in sequence space than ESM2-embedding based search (Fig 2A). While embedding-based search methods exhibit lower sensitivity in sequence retrieval compared to BLASTp and MMseqs2, they exhibit 1-3 orders of magnitude improvements in speed (Appendix C). This makes embedding-based search particularly useful for real-time search.

**Figure 2:**
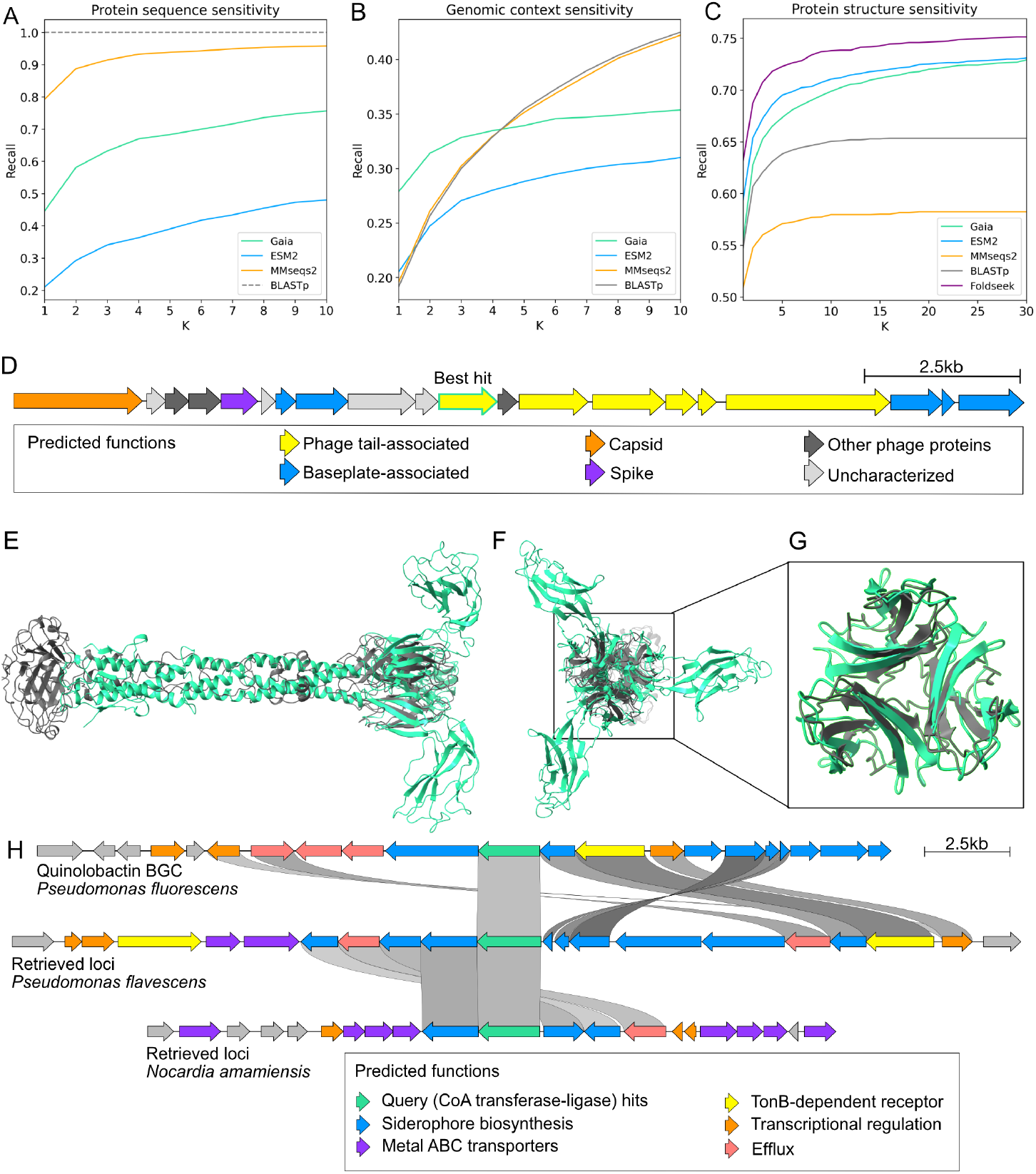
Gaia search sensitivity benchmarks and Gaia-enabled discoveries. **(A)** Protein sequence retrieval sensitivity benchmark comparing Gaia and ESM2 embedding based search (n=666), where recall (y-axis) is calculated as the fraction of searches where the best non-self BLASTp match from the OG_prot_90 was within the top K retrievals (x-axis). BLASTp is used as the ground truth (recall = 1 across all K), MMseqs2 search (with default settings) is included as a comparison. **(B)** Genomic context retrieval sensitivity benchmark comparing Gaia search and ESM2 embedding-based search (n=3,000), where recall is calculated based on the presence of a gene with a similar genomic context (at least 70% of the proteins in context matching at >50% sequence identity and >50% sequence coverage) within the top K retrievals. We benchmark these methods against MMseqs2 and BLASTp. **(C)** Protein structure-based remote homology retrieval sensitivity benchmark on the SCOPe40-test dataset (n=2,207). We compare Gaia search and ESM2 embedding-based search sensitivities in retrieving remotely homologous protein families. We benchmark against tools BLASTp, MMseqs2 and Foldseek. We show benchmarking results from other structural class levels in Appendix B. **(D)** Retrieved genomic context of the best hit phage tail protein (green highlight, center) and nearby proteins annotated as bacteriophage proteins by Gaia. **(E)** Superposed trimer structures of the query phage tail protein (green) and a phage pyosin tail fiber protein (PDB 6CU2, grey), side view. **(F)** Superposed trimer structures of the query phage tail protein (green) and PDB structure 6CU2 (grey), front view. **(G)** Aligned portion of the query phage tail protein and reference structure corresponding to the shared C-terminal lectin-like fold. The query is green, reference is gray. **(H)** Genome diagrams of the *quinolobactin* synthesis locus (top), a retrieved sequence from *Pseudomonas flavescens* (middle), and a retrieved sequence from *Nocardia amamiensis* (bottom). Retrieved sequences were obtained by using the CoA transferase-ligase *QbsK* (red) as a query.

We next compare embedding-based search’s sensitivity in retrieving proteins that are derived from similar genomic contexts. Gaia’s context-awareness enables improved retrieval of proteins that are derived from highly similar contexts (>7 out of 10 proteins in context being similar with sequence identity >50% and coverage >50%) compared to context-free ESM2 embeddings (Fig. 2B). In comparison to sequence search methods (BLASTp, MMseqs2), Gaia search exhibits improved sorting by context-similarity.

Lastly, we benchmark embedding-based search to detect remote homology using the SCOPe40-test dataset (sequences with <40% sequence identity that belong to the same structural classes) (Fig. 2C and Appendix B). Gaia search exhibits lower sensitivity for structural search compared to ESM2 embedding-based search and Foldseek, likely due to trade-offs resulting from increased sequence- and context-relevant signals in the representations. To summarize, Gaia search allows for rapid retrievals of sequences that are similar in sequence, context and structure against large and dynamic databases.

## 4 Case Studies

### 4.1 Phage tail protein

In order to demonstrate the utility of Gaia, we provide two case studies of discoveries enabled by Gaia. In our first example, we queried Gaia using a sequence with a previously uncharacterized function (Appendix D) from *Pseudomonas vancouverensis* LMG 20222. BLASTp matches of this sequence against the NCBI “nr” database returns “hypothetical proteins”. Foldseek search of the ESMFold predicted monomer structure returned proteins of various functions across databases, where lowest E-value matches with functional information were “Kinesin-like protein KIN-14F”, “Mediator of RNA polymerase II transcription subunit 21”, “uncharacterized protein” and “Helix Pomatia agglutinin with no ligands”. Using standard methods of sequence and structural search, the function of this protein is inconclusive.

With Gaia search, we first identify that the retrieved sequences occur in genomic contexts indicative of integrated prophages in bacterial genomes (Fig. 2D). Gaia’s predicted annotation for the best hit of the query protein is ‘major tail fiber protein S’, and was found to likely fold a trimer structure based on pTM and ipTM scores (ipTM = 0.68, pTM = 0.71) from AlphaFold3. Foldseek search of the trimer structure returned the lowest E-value match homology to a trimeric pyosin tail fiber protein from a *Pseudomonas* phage genome (PDB entry 6CU2, qTM score 0.17, tTM score 0.21). While the direct structural homology between these complexes is limited to a lectin-like domain near the C-termini of both proteins (Fig. 2G), both of the full trimeric structures notably form tail-like structures with highly similar length (Fig. 2E).

By comparing 100 retrieved contexts with each other, Gaia finds a diversity-generating retroelement (DGR) protein Avd (annotated as “Uncharacterized protein UU148”; not present in the best hit context shown in Fig. 2D, but frequent in other retrieved contexts) to be amongst the top five most frequently co-occurring proteins, further suggesting that this may be a DGR-associated phage tail protein. Put together, genomic context analysis validates the claim that the query protein is of viral origin and corroborates Gaia’s predicted annotation as a phage tail protein.

### 4.2 Siderophore loci

In our second example, we chose to search for putative siderophore loci. Such loci are often difficult to positively identify due to the shared similarity with biosynthetic gene clusters (BGCs) which produce products with similar chemistry, but without siderophore function. In our example, we searched for homologous genomic contexts to the Quinolobactin biosynthetic gene cluster (MIBiG accession BGC0000925) from *Pseudomonas fluorescens*. Quinolobactin is a small molecule siderophore with chemical similarity to quinolone compounds, which are used by organisms within the genus *Pseudomonas* for diverse applications including cell-cell signaling [34, 31, 19]. The Quinolobactin BGC lacks any proteins which contain the IucC condensation domain, the primary signal by which siderophores are identified in AntiSMASH [7], and so serves as an example of a siderophore locus not easily identifiable with existing BGC identification tools.

We used *QbsK* (Appendix F), a CoA transferase-ligase, from the Quinolobactin BGC as a query to Gaia. Gaia retrieved genomic contexts are putative siderophore loci across phylogenetic distances (Fig. 2F). The first example retrieval from a *Pseudomonas flavescens* genome (same genus as the query) features a distantly homologous (sequence identity 36–55%) set of siderophore synthesis-associated genes in original Quinolobactin locus. The second retrieval example from a much more distant taxon, *Nocardia amamiensis* (different phylum to the query), feature a distinctly different siderophore locus from the original query context, however, still feature siderophore-associated homologous genes with the first retrieval and other characterized siderophore loci from the genus *Pseudomonas* (Appendix G). Such homologous genes associated with siderophore synthesis include *TonB*-dependent transporters with homology to the Quinolobactin receptor, salicylate ligase and methylase proteins, and metal-specific ABC transport proteins. The full annotations for selected putative siderophore loci identified by this method are viewable in Appendix H.

These putative synthesis loci identified using Gaia search could not be identified using HMM rule-based methods such as AntiSMASH, nor with sequence or structural search, as the query protein function (CoA transferase-ligase) is ubiquitous and non-specific to siderophore synthesis. Through this example, we demonstrate Gaia’s utility for discovering novel potential BGCs that share similar genomic architectures.

## 5 Conclusion

Gaia provides a unique capability to bioinformatics researchers by incorporating information about the genomic context of proteins. While BLASTp and other sequence alignments approaches excel at finding primary sequence similarities and Foldseek focuses on structural relationships, Gaia provides the ability to identify related proteins with conserved genomic contexts with search speed orders of magnitude faster than existing methods. This capability enables researchers to address complex questions in comparative genomics, such as identifying conserved gene clusters across distantly related organisms, detecting horizontal gene transfer events, and elucidating the function of hypothetical proteins based on their genomic neighborhood. By integrating these approaches, researchers can gain a more comprehensive understanding of protein function and evolution in genomic data, particularly in scenarios where sequence or structural similarity alone may be insufficient.

## Methods

### gLM2 finetuning

We finetune the pre-trained gLM2 model for the retrieval task with a 2-stage approach, similar to how text embedding models are finetuned from pre-trained BERT models [36, 46]. In the first stage, we finetune gLM2_650M on Uniref50 for one epoch. We use a learning-rate of 1e-4 with cosine decay and batch size of 256. This reduces train-test mismatch as gLM2 is pre-trained on genomic contigs instead of individual proteins.

In the second stage, we align representations with protein structure, by training a linear projection layer on mean-pooled, frozen gLM2 representations. We use 2.3M structural clusters from the AlphaFold Database [5], and train the linear layer to maximize the cosine-similarity between proteins in the same cluster. During training, we randomly sample pairs of protein sequences with the same cluster assignment, and optimize the InfoNCE contrastive loss [50]. We use a large batch size of 32,768 to increase the number of in-batch negatives, and train for 30,000 steps with learning-rate 1e-4. We use representations from the middle layer of gLM2 (16th layer), as previous studies have shown improved transfer-learning ability compared to the final layer [39, 53]. The output dimensionality of the projection layer is 2.5x smaller than the gLM2 hidden dimension (512 vs 1280 dimensions), improving the scalability of vector search across large databases.

### Database creation

All protein coding sequences from the OpenGenome dataset [12] consisting of 131,744 prokaryotic and viral genomes available on INSDC databases (https://www.insdc.org), (Cochrane et al. 2016) and retrieved from IMG/M (Chen et al. 2023)were extracted and clustered at 90% sequence similarity using MMseqs2 (commit 16e46) [45] cluster with configuration –min-seq-id 0.9 -c 0.9 yielding the OG_prot_90 database consisting of 85,007,726 centroid sequences.

### Vector database search

Gaia’s search uses the Qdrant vector database library [37] for rapid similarity search against the embeddings database. A query protein sequence is first embedded with the gLM2 model, and the resulting vector is used to search for similar sequences in the database using the cosine-similarity metric. We use the Hierarchical Navigable Small-World (HNSW) algorithm [30] to efficiently search for approximate nearest-neighbors across millions of protein embeddings.

### Gaia output visualization

The output includes a UMAP [32] representation of the retrieved vector sequences, colored optionally by cosine similarity or by BLASTp percent identity to the input vector. Additionally, an ESMfold-generated protein structure is displayed, along with optional overlays of detected Pfam HMM domains on the query protein. Genome context visualization diagrams, BLASTp identity and coverage, an ESMfold-predicted structure, and a Needleman-Wunsch [35, 11] alignment between the query and each returned subject sequence is available in an expandable accordion for each of the 100 returned subject sequences. HMM annotation in Gaia is performed using pyHMMER [25] by running the Pfam HMM database (v37.0) with the gathering (GA) precomputed bitscore thresholds against the query and subject sequences. Predicted protein structures for query and subject sequences are generated using ESMFold and visualized with Molstar [42]. Coverage values and percent identity were generated with BLASTp using the ncbi-blast+ suite [10]. Needleman-Wunsch alignment was generated using Biopython [11]. Top five most frequently co-occurring proteins were calculated by clustering all retrieved context protein embeddings with DBSCAN [15] with cosine metric and eps=0.05. The Gaia frontend was generated using the Nitro UI software package [22].

### Functional annotation

Functional annotations for retrieved sequences are generated by aligning pLM representations with functional text-annotations. We train a CLIP-like model [38] to align ESM2 representations with text annotations, encoded using a pre-trained text encoder dmis-lab/biobert-v1.1 [26]. The model is trained on the SWISS-prot database for 5 epochs with a batch size of 6,000 protein to text pairs, and learning rate of 1e-4. We keep the ESM2 model frozen, and train only the projection layer and the text encoder. We precompute functional annotations for each protein in OG_prot_90 by finding the closest SWISS-prot protein embedding in cosine-similarity.

### Sequence retrieval sensitivity benchmark

A random set of 1200 sequences was selected from OG_prot_90. Sequences were then clustered using CD-HIT 4.8.1 [18] with the following parameters: -c 0.5 -n 2. The resulting clustered sequences were then used as a query to BLASTp v2.12.0+, and searched against a BLAST database of OG_prot_90. Only 666 sequences with a non-self hit with thresholds 75% identity and 70% reciprocal coverage were kept. Recall was calculated by determining whether the best non-self BLASTp hit is included in the first K retrievals using embedding-based search. For the MMseqs2 baseline, we ran the easy-search pipeline with default parameters.

### Genomic context sensitivity benchmark

A random set of 3000 proteins were selected from OG_prot_90 using the seqkit sample function [43]. We then queried our qdrant database to compare the query protein’s genomic context, where context is defined as up to five protein coding genes upstream and five protein coding genes downstream in the genome. We use BLASTp to calculate the homologous fraction of the surrounding proteins between the query and the top ten retrieved examples, where two proteins are considered homologous if there is sequence identity >50% with coverage >50%. If more than seven out of ten genes in the retrieved protein’s context are homologous to those in the query protein’s context, we consider the retrieval to be a correct retrieval. For the BLASTp and MMseqs2 baselines, we searched the same 3000 proteins against the OG_prot_90 database using default parameters and compared the homologous fractions of the genomic contexts of top 10 non-self hit proteins using the same parameters as above.

### SCOPe-40 retrieval sensitivity benchmark

The SCOPe-40 2.01 [17] test set is used to benchmark structural sensitivity, which includes 2,207 protein sequences with family, superfamily and fold structural classification. For each protein sequence, we embed using the gLM2_embed model, and retrieve the closest 30 proteins in the test set (excluding self-hit) with the cosine similarity metric. We compute recall by checking if any of the top-k retrievals match the query’s structural family. The ESM2 baseline is computed using the middle layer hidden representation from the ESM2 650M model. All other baseline method hits are downloaded from https://wwwuser.gwdguser.de/~compbiol/foldseek/scop.benchmark.result.tar.gz.

### Phage tail protein case study methods

Predicted protein structures for Figure 2 were generated using AlphaFold 3 [1]. Structure search was performed using the Foldseek-Multimer web service [24]. The protein reference structure used was obtained from RCSB-PDB [6, 40]. The genome context diagram in Figure 2D was generated using clinker [20], from IMG genome ID 2667527228, contig ID Ga0074689_11, part of predicted prophage IMGVR_UViG_2667527228_000002. Siderophore loci case study methods Genome context diagrams were generated using clinker. Reference BGC sequences for loci producing the siderophores quinolobactin (BGC0000925), yersiniabactin (BGC0002570), enterobactin (BGC0000343), and pyochelin (BGC0000412), including their annotations, were obtained from MIBIG [48].

### Siderophore loci case study methods

Genome context diagrams were generated using clinker. Reference BGC sequences for loci producing the siderophores quinolobactin (BGC0000925), yersiniabactin (BGC0002570), enterobactin (BGC0000343), and pyochelin (BGC0000412), including their annotations, were obtained from MIBIG [48].

## Data and model availability

gLM2_embed is available at https://huggingface.co/tattabio/gLM2_650M_embed. OG_prot_90 is available at https://huggingface.co/datasets/tattabio/OG_prot90. All benchmarking code is available at https://github.com/TattaBio/gaia-benchmark.

## Acknowledgements

The work conducted by the US Department of Energy Joint Genome Institute (https://ror.org/04xm1d337) is supported by the US Department of Energy Office of Science user facilities, operated under contract no. DE-AC02-05CH11231. S.R. was supported by the U.S. Department of Energy, Office of Science, Biological and Environmental Research, Early Career Research Program awarded under UC-DOE Prime Contract DE-AC02-05CH11231. We thank Milot Mirdita, Lingdong Shi, Jordan Hoff, Brooke Travis, Luis Valentin Alvarado, Mark Little, Juan Pierella Karlusich, Luis Valentin Alvarado and Matthew Schechter for participating in our user interviews and providing us feedback.

## Appendix A Additional features of Gaia web server

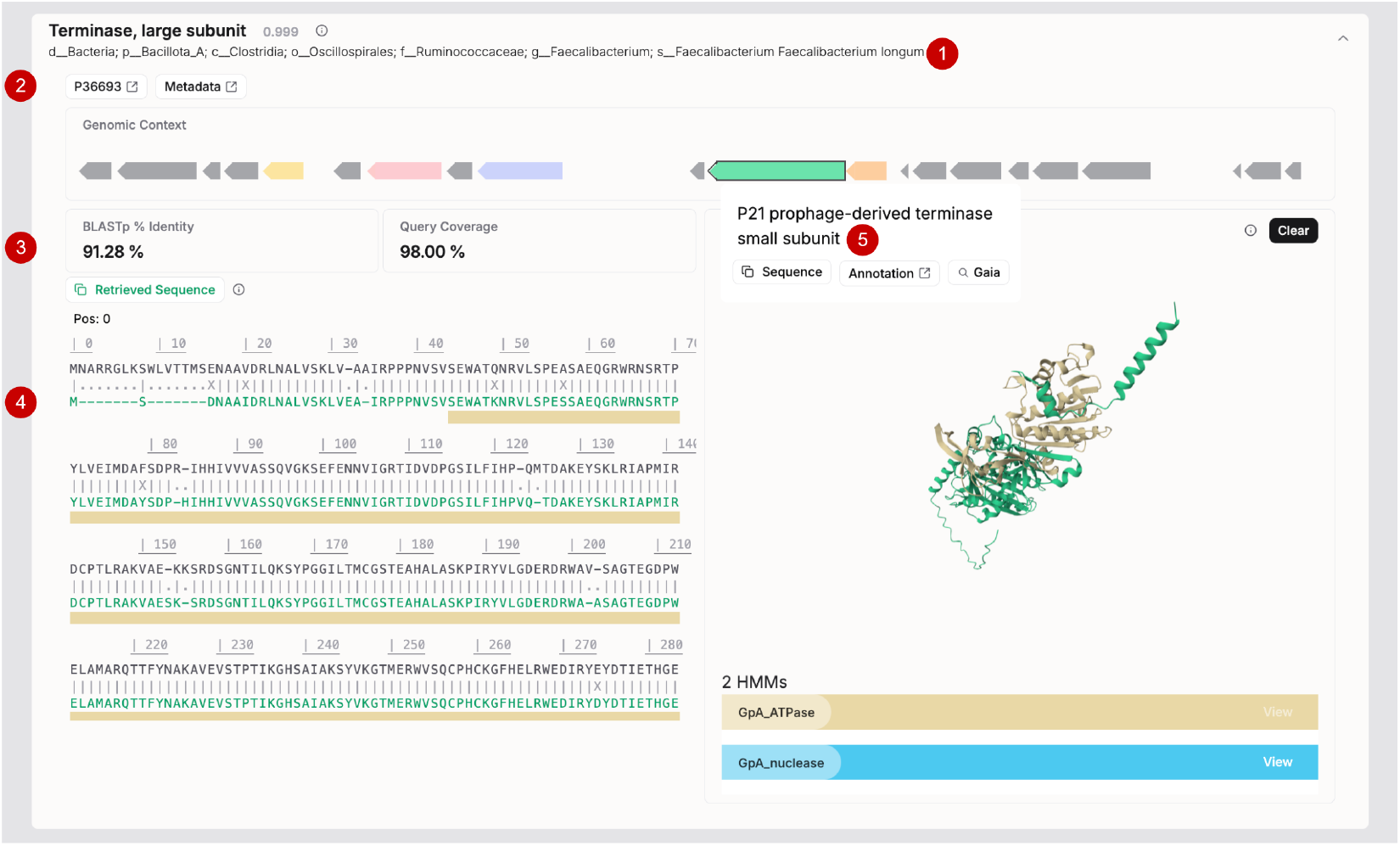

**An open accordion view of a search result. (1)** Predicted Swiss-Prot annotation with level of confidence (in gray). The full GTDB style taxonomy of the genome this sequence is found is shown below the annotation. **(2)** Links to the Swiss-Prot annotation this sequence was matched to and metadata information of sample this sequence was found in. **(3)** BLASTp calculation between the retrieved sequence and the query. **(4)** Global alignment (Needleman-Wunsch) of the retrieved sequence against the query. **(5)** Protein-coding genes found in the same context are annotated and can be iteratively searched upon clicking.

## Appendix B SCOPe40-test remote homology benchmark at different structural class levels (Fold and Superfamily)

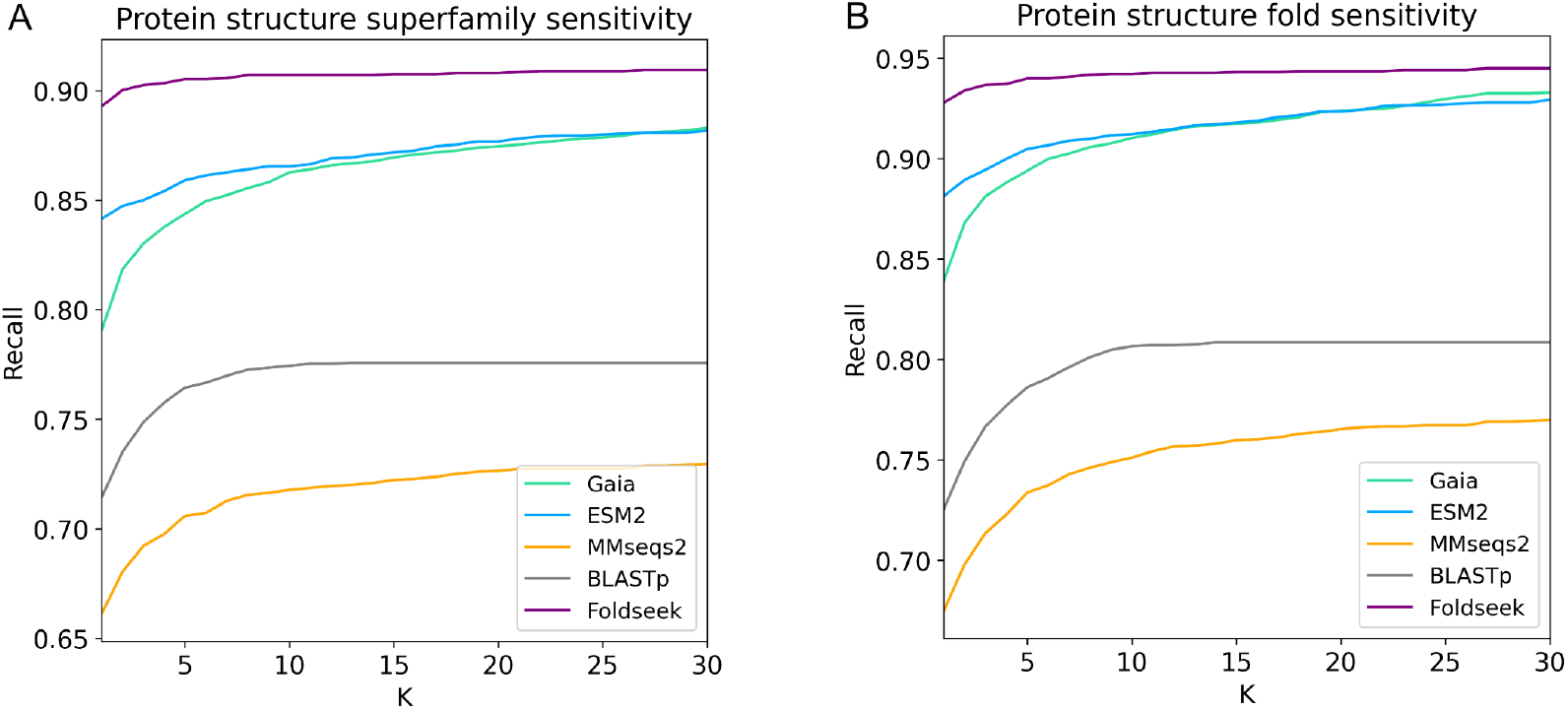

## Appendix C Speed benchmarking

| # Sequences | BLASTp | MMseqs2 | Gaia |
| --- | --- | --- | --- |
| 1 | 18.7 | 8.3 | **0.14** |
| 10 | 98.2 | 15.3 | **0.2** |
| 100 | 1070 | 21.3 | **1.3** |
| 1000 | 10,768 | 32.5 | **14** |

We compare embedding-based search methods against BLASTp and MMseqs2 for searching 1, 10, 100, 1000 sequences against OG_prot_90. Target database (OG_prot_90) creation times are not included in this calculation. BLASTp was run with the default settings using 20 threads. MMseqs2 was run with default settings with index in memory (`–db-load-mode` 2 after `touchdb`) and 20 threads. Gaia includes GPU inference time for sequence embedding (T4 GPU) and vector search using Qdrant with default HNSW parameters (`m=16, ef_construct=100`) and 20 threads, with k=100 nearest neighbors. Time is reported in seconds.

## Appendix D Putative Phage Protein Sequence

~~~
MANLNGATGVAAIFKAFLRKLETTDPKHPDTWNPNYQTLIDNDVFLKAFADEVSTARGSQPSLKDRLVAIEQT
QASLSPEYIDELTAAVKYALDQAGVANRSIRALKSQLQQEGELLIENRGIVSGCTATKSTTAARNLNLAAGVC
FANGRAYSVDSGNNMASVPSNISAGNASAVVYLYRSGNGWKMAVTAIGEAVPAGAIRLYNVTIPPNSTDATDP
TLANVTLTSVRRVEVGFPQYLDTPVSQFVAINNLSANDFRLDFEVVSAEGAPCERKSLSVPSRATNGFTLELA
SAADNVLVRYRVSKLNN
~~~

## Appendix E Full predicted annotations for the genomic context of putative phage tail protein

Annotations for the genomic context of the putative phage tail protein highlighted in Figure 2 are provided at https://doi.org/10.5281/zenodo.14182698.

## Appendix F QbsK protein sequence used as query for siderophore loci discovery

~~~
MSLLKHLTLHLSASAWSAELAPVAEALARRLKAQGGQVATGGQTLCDDAGHQVQLHLRPWPAPSTVAASAALV
EAVVGLTALHQRSSGEPLPLGVDYCATFTASLLLTAALASLLGQARGLPAARLAMSHGGAALLAIGQYLAMDS
ADGGYPPEPAPPADAVRPPFTSADGVVFELEALNPDAWLRFWQQAGVDIAVAGKGWRPFMQRYTRATAWLPAA
LMRAASEHDFAQLQAMAKEAGTALCALQRWQDCRNQPVFQPWIQSGGWRCTEFAPGPGVNGPDEQQPPQHAGL
PLQGITVVECCRLIQGPLAGHVLRLLGATVIKVEPPGGDPMRGMPPMAGEISAHFDAINRSKQTVQIDLKAPT
GRAELLALLAHADVFLHNWAPGRDEQLALQPHTLARLLPRLIYVSASGTGPAPTSDQPLGTDFMIQAYSGLAE
RIRQPHGAAGGALITLVDVLGAVAAAEGIVAALYARQRDGRGRYLDSCMAGAAALLLAGQPDGQAPEPAEAFA
CRDGWLMLDRAQSARGRDGLARLQADPGLAPWCAEQDTHACCRALQAMGVMAVAVTADLRQLPDLPLLAGALR
QSAYLHVNNPWEISNL
~~~

## Appendix G Siderophore genome contexts with reference siderophore loci from the genus *Pseudomonas*

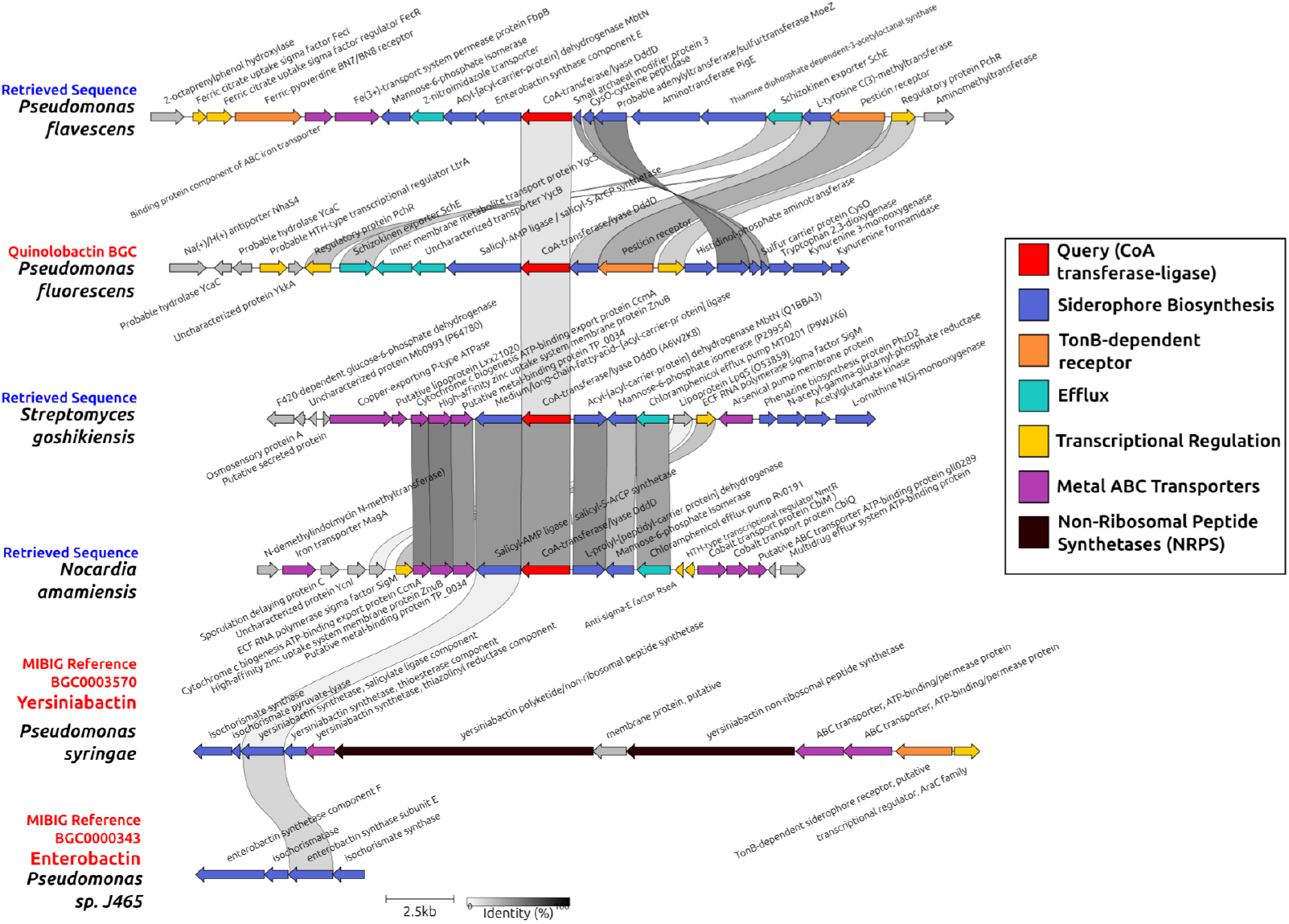

Three retrieved putative siderophore producing loci obtained from searching with Gaia (blue) and three reference siderophore-producing loci from the genus *Pseudomonas* (red). Retrieved sequences were obtained by using Quinolone CoA transferase-ligase QbsK as a query to Gaia. Proteins homologous to the salicylate ligase subunits of Enterobactin and Yersiniabactin synthase were present in all analyzed genome contexts. Putative siderophore producing function in the retrieved sequences is supported by the presence of siderophore receptor-like TonB-dependent receptor genes, metal transport machinery, and efflux pumps with homology to the Schizokinen siderophore efflux system.

## Appendix H Full annotations for selected putative siderophore loci

PFAM annotations, Gaia annotations for retrieved sequences, and MIBIG annotations for reference sequences for the genomic contexts of retrieved putative siderophore producing loci and reference sequences from MiBIG are provided at https://doi.org/10.5281/zenodo.14182698.

